# Syntaxin-1 and syntaxin binding protein 1-1 are required for planarian behavior and brain regeneration

**DOI:** 10.64898/2026.10.05.756827

**Authors:** Kendall B. Clay, Pierce A. Roberts, Karl F. Lechtreck, Rachel H. Roberts-Galbraith

## Abstract

**Background:** Nervous system regeneration requires creation of new neurons and their integration into new and existing networks. How new neurons find their targets and reconnect in the environment of regenerating tissue remains a major gap in knowledge. The planarian *Schmidtea mediterranea* is well suited to the study of neural regeneration due to the remarkable ability of planarians to regenerate their entire body, including the nervous system. However, we have not yet been able to explore synaptogenesis in regeneration using planarians due to a lack of tools to study synapses. Here we investigate potential players in planarian synapse, focusing on the conserved syntaxin protein family, which functions in neurotransmitter vesicle fusion at the synapse.

**Results:** We found that reduction of either *syntaxin-1* (*stx1*) or *syntaxin-binding protein 1-1* (*stxbp1-1*) decreased locomotion and impeded brain regeneration, specifically in synapse-rich regions.

**Conclusions:** Our observations support STX-1 and STXBP1-1 as bona fide synapse components, providing a starting point for studies neuronal connectivity during homeostatic remodeling and regeneration.

## Introduction

Neuronal connectivity occurs through signaling contacts between neurons known as synapses. Synapses involve diverse protein components, including cell-cell adhesion molecules, trafficking machinery, transporters, receptors, and structural proteins (e.g. cytoskeletal regulators). Chemical signaling between neurons requires transport of vesicles down the axon and then, upon stimulation, fusion of neurotransmitter-filled vesicles for their release into the synaptic space. The process of vesicle fusion and neurotransmitter release itself requires a host of proteins, including synaptotagmin, synaptobrevin, syntaxin, and syntaxin-binding proteins, which work together as part of a larger Soluble N-ethylmaleimide-sensitive factor Attachment protein Receptor (SNARE) complex^1^ (Fig. 1A). Components of the SNARE complex are present within the synapse on vesicles and on the cytoplasmic side of the pre-synaptic cell membrane, where they interact to facilitate proper neurotransmission^2^. Though synapses have been well characterized molecularly and functionally in several animal species, formation and function of synapses has been best understood in the context of embryonic development and homeostasis. Much less is known about how synapses are reestablished in a regenerative context (regenerative synaptogenesis) and how injury may impact synapse form and function.

**Figure 1:**
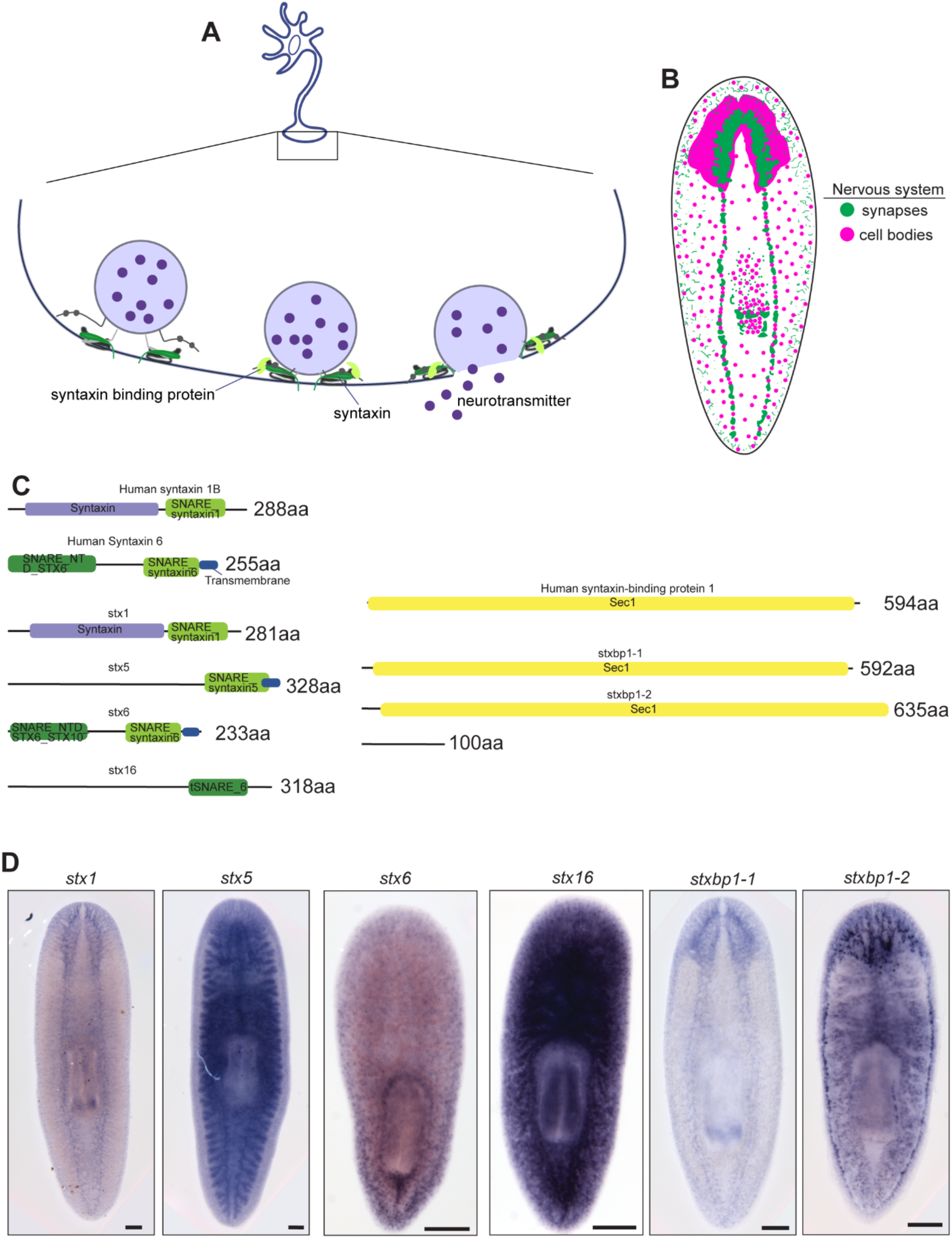
Planarians have four *syntaxin* homologs and two *syntaxin-binding protein* homologs. A) Diagram depicting a presynaptic neuron terminal. Vesicles of dark purple neurotransmitters are shown fusing with the membrane with the help of SNARE complex proteins (gray), syntaxin (green), and syntaxin-binding protein (light green). B) Nervous system schematic depicting a planarian nervous system made of cell bodies (magenta) and synapses (green). C) Domain architecture for human and planarian STX and STXBP proteins. D) Colorimetric ISH showing expression patterns of planarian *stx* and *stxbp* homologs. Scale bar = 200 µm.

Regeneration, defined as the ability to regrow tissue *de novo*, occurs to varying degrees across the animal kingdom (for review, see ^3^). Although wound-healing of structures like the epidermis is widespread, few organisms can robustly regenerate their entire nervous system^4–6^. One organism capable of whole brain regeneration is the planarian^7^. Planarians regenerate all organs and mature cell types thanks to their adult pluripotent stem cells^8–11^, which constantly divide to replenish the stem cell pool and to replace mature cell types of the body after injury and during homeostatic turnover^12,13^. The nervous system includes a central nervous system (bilobed brain and ventral nerve cords), peripheral nervous system, and pharyngeal nervous system^14^ (Fig. 1B). Planarians complete full brain regeneration in just seven days after head amputation^15^, with restoration of dozens of neuronal and glial cell types in the right ratios and locations^16–19^. Synapses, marked with anti-synapsin, also form quickly^20,21^. Animal behavior—including movement toward food and away from light^22–26^—also resumes post-injury, indicating functional restoration and connectivity. However, we do not know how new planarian brain tissue is rewired: how new neurons find their targets; how new synapses are created and matured; and how connections are remodeled during brain regeneration. These gaps in knowledge persist partly because planarian synapses have not yet been characterized. Further, understanding the planarian synapse could reveal whether and how neuronal signaling promotes regeneration.

In other animals, neurotransmitter release at the synapse is coordinated through protein families that mediate docking of neurotransmitter-laden vesicles and membrane fusion^1^. Two key players are SNARE protein syntaxin and syntaxin binding protein (which is also called UNC-18/munc18), which cooperate to mediate exocytosis^2,27–29^. We reason that because syntaxins have been shown to be key molecules in neurotransmission in other organisms they may be involved in planarian neurotransmission^30–33^. Importantly, though, SNARE machinery is also involved in non-neuronal exocytosis^34–37^. Therefore, we must start with a list of putative synaptic proteins and identify those most likely to be *bona fide* neuronal synapse components through localization and functional studies. In this work, we identified four homologs of *syntaxin* and two homologs of *syntaxin binding protein* in planarians. We found that *syntaxin 1* (*stx1*) and *syntaxin binding protein 1-1* (*stxbp1-1*) are present throughout the nervous system, are required for movement and response to stimuli, and are required for proper regeneration of the synapse-rich neuropil in planarians. We conclude that STX1 and STXBP1-1 are conserved synapse proteins in planarians.

## Results

We reasoned that the genes important for planarian chemical synapses should meet two criteria. First, they should be expressed in neural tissues, with protein present at the synapse itself. Second, synapse proteins should promote neural signaling, meaning that disrupting a *bona fide* synaptic component would likely disrupt some aspect of animal behavior. To identify synapse factors in planarians, we began by using BLAST similarity to identify planarian homologs of *syntaxin* and *syntaxin binding protein*. We narrowed down our list via reciprocal BLAST and focused on matches with e-values smaller than 1*10^-20^. We identified four homologs of syntaxin and two homologs of syntaxin binding protein meeting this threshold. We also used phylogenetic analyses to name each homolog (Supp. Fig. 1A-B). We then confirmed conserved protein domains for all homologs (Fig 1C). To identify which homologs were enriched in neurons, we assessed expression patterns for each gene using colorimetric *in situ* hybridization (ISH). We found that *stx1* and *stxbp1-1* were expressed throughout the central nervous system in the brain and ventral nerve cords, and that *stx5* was expressed faintly in the brain as well as in the intestine (Fig. 1D). *stx6*, *stx16*, and *stxbp1-2* were each expressed in the peripheral nervous system (Fig. 1D). Several of these genes, such as *stx5*, *stx16*, and *stxbp1-2*, were also expressed in non-neural tissues which was clear in both ISH and available transcriptional atlases (^38^; Fig 1D). We concluded that *stx1* and *stxbp1-1* fit the first synapse criterion of neuronal expression.

To understand gene function and test our second prediction for synapse protein-encoding genes, we knocked down the *syntaxin* and *syntaxin binding protein* genes individually through RNAi (Fig. 2A). During the RNAi paradigm, we began to notice differences in movement after *stx1*(RNAi) or *stxbp1-1*(RNAi) that warranted further investigation. We saw slower gliding movement as well as occasional inch-worming behavior in RNAi animals, which can be seen in still images as a transient shortening of the body (Fig. 2B-C). To confirm differences in movement, we placed animals in the center of a well-lit dish and assessed whether they moved out of this region of the dish after one minute. Only 2/12 controls stayed in the center of the dish, but 12/12 of the *stx1*(RNAi) and 12/12 of the *stxbp1-1*(RNAi) worms stayed in the center (Fig. 2D). We also noted that *stx1*(RNAi) and *stxbp1-1*(RNAi) animals exhibited an unusual, elongated posture which we quantified using a length-to-width ratio (Fig. 2E-F). Because of these movement and posture differences, we suspected there may be a defect in cilia-mediated and/or muscle-mediated movement. This would support our hypothesis of a role for *stx1* and *stxbp1-1* in neuron function, as neurons control both muscle-mediated and cilia-mediated movement^39–41^. Because gliding motion is driven by the rhythmic beating of motile cilia on the ventral epidermis, we assessed a role for each gene in cilia-mediated movement. Ciliary length and beat frequency were similar in control animals and *stx1*(RNAi) or *stxbp1-1*(RNAi animals) (Supp. Fig. 2A-B). In control animals, we also observed metachronal waves, indicative of tissue-wide ciliary coordination (Fig. 2G-H). In contrast, cilia beating was more uncoordinated in the *stxbp1-1*(RNAi) and *stx1*(RNAi) animals, offering a possible explanation for the reduction in gliding velocity (Fig. 2G-H).

**Figure 2:**
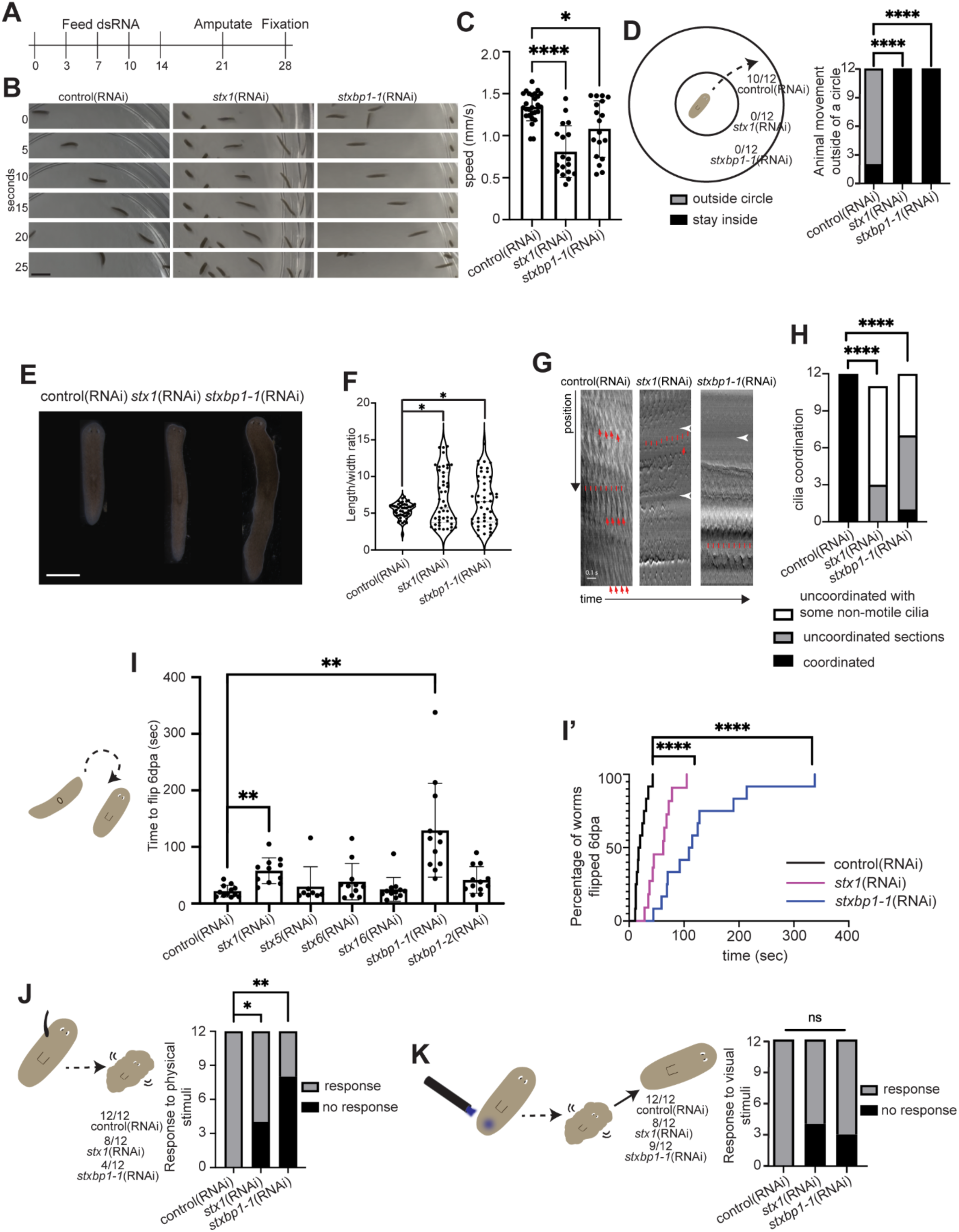
Planarian *stx1* and *stxbp1-1* are required for planarian behaviors, especially after regeneration. A) RNAi paradigm. B) Live images showing animal position over time in knockdown versus control animals. C) Speed in knockdowns versus controls in mm/s. Differences were evaluated with a Brown-Forsythe and Welch ANOVA with Dunnett’s T3 multiple comparisons. D) Schematic of movement assay performed post-amputation. Numbers show the fraction of animals to move from the inner circle after 1 minute. Differences were assessed using Fisher’s exact test, adjusted for multiple comparisons. E) Live images showing differences in body shape and posture. F) Length/width ratios for control and knockdown animals as shown in E. Differences were evaluated with a Brown-Forsythe and Welch One-way ANOVA with Dunnett’s T3 multiple comparisons. G) Example kymograms from control(RNAi), *stx1*(RNAi), and *stxbp1-1*(RNAi) animals used to analyze coordination and measure beat frequency. White arrowheads point to regions of the kymograph devoid of motile cilia. Red arrows point to regions of metachrony where several cilia beat together, resulting in long parallel lines. H) Representative kymograms from the anterior head of each animal (N = 11-12 animals per condition with 12 kymograms/animal) were used to score coordination. Kymograms with long parallel lines down the length of the image were scored as coordinated. Kymograms with several different beat patterns were scored as uncoordinated sections. Kymograms with sections of non-motile cilia were scored as uncoordinated with some non-motile cilia. Assessment of cilia coordination for 11-12 animals from each condition, with Fisher’s exact test, adjusted for multiple comparisons. I) Righting assay depicting time to flip over at 6 dpa, with differences assessed using a one-way ANOVA with Dunnett’s multiple comparisons test. I’) Data from the righting assay graphed as percentages with the differences assessed using a Mantel Cox log-rank test. J) Schematic showing the fraction of worms responding to touch stimulus, evaluated using Fisher’s exact test, adjusted for multiple comparisons. K) Schematic showing the fraction of worms responding to visual stimulus, with differences assessed using Fisher’s exact test, adjusted for multiple comparisons. Scale bar = 5 mm (B), 1000 µm (D). ns = not significant, * P ≤ 0.05, ** P ≤0.01, *** P ≤0.001, **** P ≤0.0001.

Previously, our lab established a righting assay that can be used to assess coordinated muscle-based movement^7,42^. We completed a righting assay in which we placed animals ventral side up and assessed the time it took for them to flip over. After amputation and regeneration, *stx1*(RNAi) and *stxbp1-1*(RNAi) animals took much longer to right themselves (Fig. 2I-I’). The impact of RNAi was much milder before amputation and regeneration (Supp. Fig. 2C-C’). We conclude that *stx1*(RNAi) and *stxbp1-1*(RNAi) do impact muscle movement and coordination during righting. In summary, our observations support a role for *stx1* and *stxbp1-1* in neuron function, as they are required for both muscle-mediated and cilia-mediated movement that is under neuronal control^39–41^.

Another role for the nervous system is in responding to stimuli. To assess response to physical and visible stimuli, we assessed response to touch and bright light. We first used an eyebrow hair to touch animals in their trunk regions and saw that although all 12/12 controls responded, only 8/12 *stx1*(RNAi) and only 4/12 *stxbp1-1*(RNAi) responded to touch with contraction (Fig. 2J). Next, we used a visual stimulus, blue laser light, because planarians are averse to shorter wavelengths of light^43^. Again, we saw that all 12/12 controls responded to bright light but only 8/12 *stx1*(RNAi) and 9/12 *stxbp1-1*(RNAi) responded by contracting or moving (Fig. 2K). In conclusion, abrogating *stx-1* and *stxbp1-1* function caused stimulus response after brain regeneration, though whether the defect is in sensory function itself and/or movement after stimulus cannot yet be determined. Taken together, our evaluation of gliding, righting, and stimulus response behaviors indicate that STX1 and STXBP1-1 fulfill the second criterion for genes that potentially function in the planarian synapse.

Because we saw similar behavioral effects in both knockdowns, we wanted to assess whether *stx1* and *stxbp1-1* could work together in the same process. To assess potential cooperation, we completed a double knockdown of these genes and did not see exacerbation of RNAi phenotypes prior to amputation and regeneration (Supp. Fig. 2D-D’). However, after regeneration, the effect of RNAi was additive, with half of double knockdown animals failing to right themselves after 6 minutes (Fig. 3A-A’). To support the hypothesis that these proteins cooperate, we found that *stx1* and *stxbp1-1* mRNA were co-expressed 50% of the time (Fig. 3B). Additionally, we verified that *stxbp1-1* is indeed nearly exclusively expressed in neurons via co-staining with a pool of neural probes designed to label the majority of neurons (^38^; Fig. 3C).

**Figure 3:**
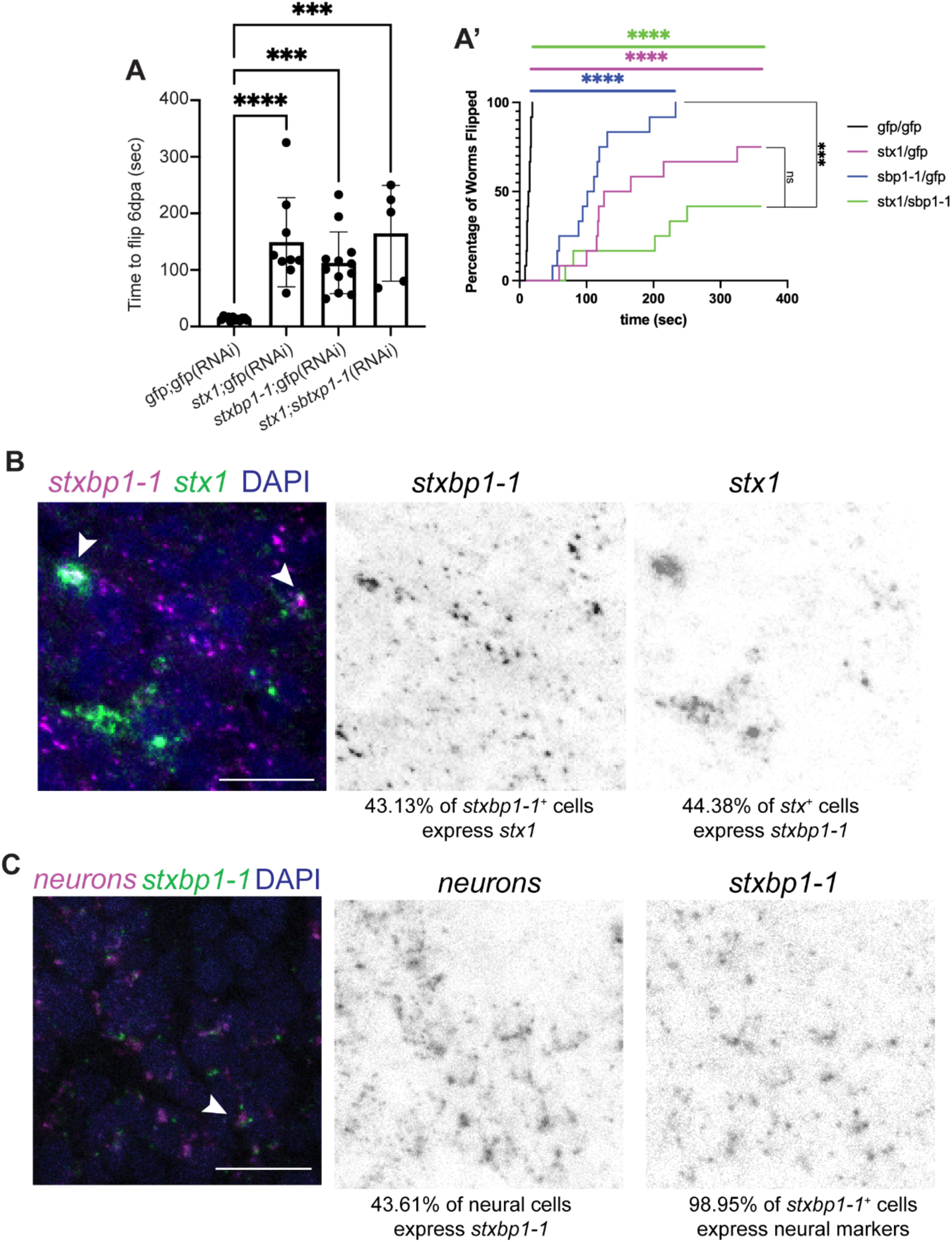
Planarian *stx1* and *stxbp1-1* cooperate in a subset of neurons. A) Double knockdown experiments showing average time to right after 6 dpa. Differences were assessed with one-way ANOVA with Dunnett’s multiple comparisons test. A’) Graph showing the percentage of animals righted over time, with differences assessed using a Mantel-Cox log rank test. Double knockdown animals fail to right more frequently than *stx1/gfp* knockdown animals (P< 0.001) but were not significantly different from *stxbp1-1/gfp* animals. B) Fluorescent ISH showing co-expression of *stx1* (magenta) and *stxbp1-1* (green) with DAPI (nuclei – blue). In each single channel, the percentage of cells co-expressing the other marker is written below. C) Fluorescent ISH showing co-expression of a pool of neural markers [*synapsin* (*dd_Smed_v6_3135_0_1*)*, synaptotagmin* transcripts (*dd_Smed_v6_4222_0_1, dd_Smed_v6_6730_0_1,* and *dd_Smed_v6_6920_0_1*)^38^; magenta] and *stxbp1-1* (green) with DAPI (nuclei – blue). In each single channel, the percentage of cells co-expressing the other marker is written below. Scale bar = 20 µm.

Ultimately, one goal of this project was to identify factors of the planarian neuronal synapse. A second goal was to assess whether and how perturbation of neurotransmitter release affects regeneration. We therefore knocked down homologs of *syntaxin* and *syntaxin binding protein*, amputated heads, waited 7 days for head regeneration, and then fixed the animals for *in situ* hybridization (ISH). If perturbing neurotransmitter release affects regeneration, we expected to see reduced brain size after regeneration in *stx1*(RNAi) and/or *stxbp1-1*(RNAi) animals. We used the marker *choline acetyltransferase* (*ChAT*) and calculated brain-to-body ratios after regeneration (Fig. 4A). Indeed, we found that *stx1*(RNAi) or *stxbp1-1*(RNAi) led to smaller brains after regeneration whereas *stx5*(RNAi) led to lysis and rare survivors with larger brains after regeneration (Fig. 4B). We note that brain regeneration was not additively impaired by simultaneous knockdown of *stx1* and *stxbp1-1,* although variance in the double knockdown condition was increased (Supp. Fig. 2E-F).

**Figure 4:**
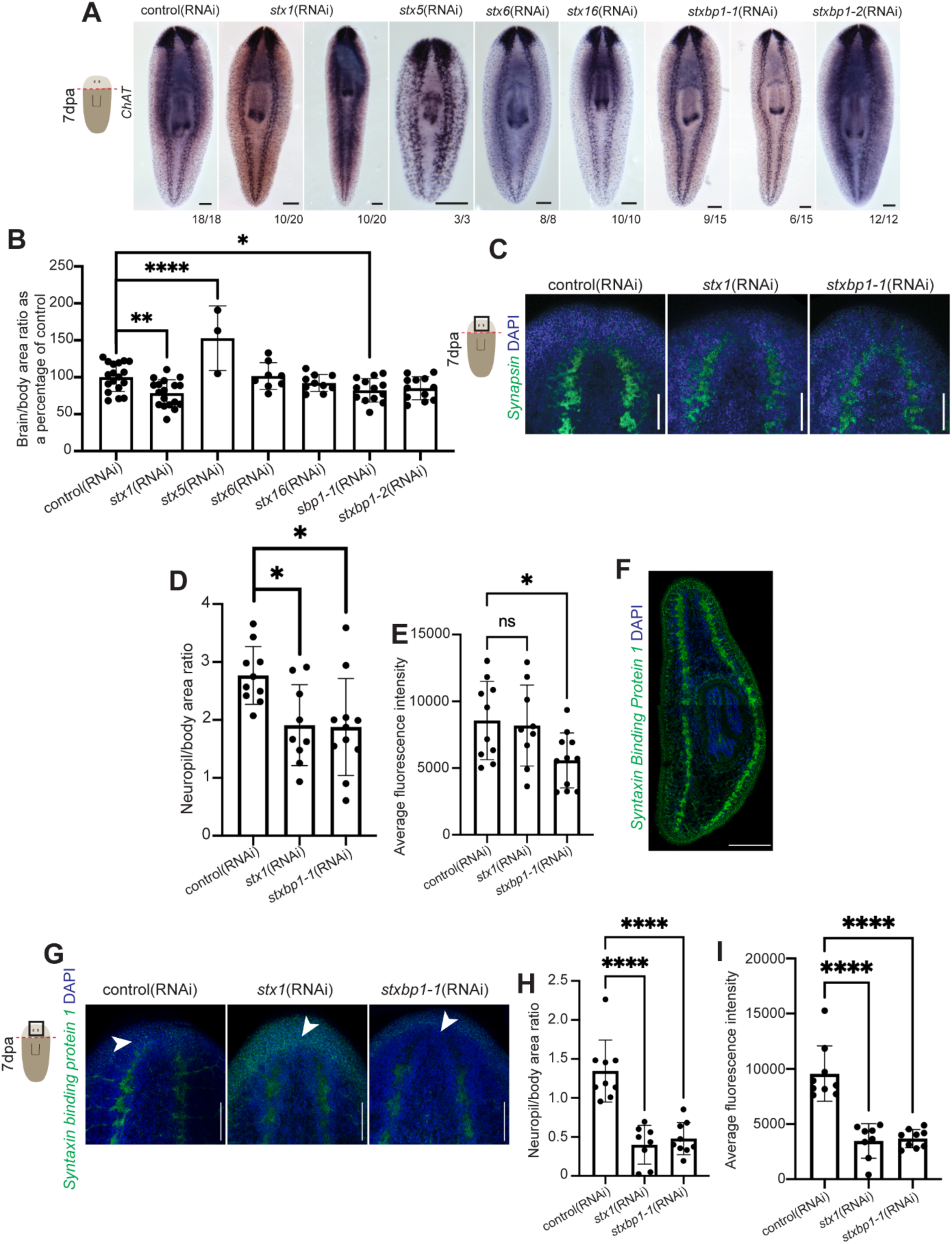
Knockdown of *stx1* or *stxbp1-1* causes a reduction in brain regeneration through reducing synapse volume. A) Colorimetric ISH depicting *ChAT* expression after knockdowns showing either 1 or 2 representative images. B) Quantification of brain-to-body ratio from A. One-way ANOVA with Dunnett’s multiple comparisons test. C) Fluorescent immunofluorescent images depicting anti-synapsin (green) and DAPI (nuclei – blue) in the regenerating head. D-E) Quantification of C depicting neuropil-to-body ratio (D) and average fluorescence intensity (E). One-way ANOVA with Dunnett’s multiple comparisons test. F) Fluorescent immunofluorescent tiled image depicting anti-syntaxin binding protein 1 (green) and DAPI (nuclei – blue) in the whole planarian body. G) Immunofluorescence images depicting syntaxin binding protein 1-1(green) and DAPI (nuclei – blue) in the regenerating head with the neuropil denoted by white arrowheads. H-I) Quantification of G depicting neuropil-to-body ratio (H) and average fluorescence intensity (I). One-way ANOVA with Dunnett’s multiple comparisons test. Scale bar = 200 µm (A, F), 100 µm (C, G), 50 µm (G). ns = not significant, * P ≤0.05, ** P ≤0.01, *** P ≤0.001, **** P ≤ 0.0001.

We were surprised to see that *stx5*(RNAi) caused larger brains but noted that there were only 3 surviving animals in the experiment in Fig. 4A-B because *stx5*(RNAi) causes lesions, lysis, and death (Supp. Fig. 3A-B). We thus completed an alternative paradigm with early amputation and were surprised to again find that surviving *stx5*(RNAi) animals had significantly larger brains than controls (Supp. Fig 3C-E). In an RNAi paradigm with no amputation, we observed that about half of *stx5*(RNAi) animals underwent head regression^44^ and therefore had smaller brain size, but the animals that avoided head regression had significantly larger than normal brains, showing that the impact of Stx5 on brain size is not dependent on regeneration (Supp. Fig 3F-G). Interestingly, *stx5* is required for both exocytosis and inter-organellar trafficking, including vesicle trafficking between the Golgi and endoplasmic reticulum^45,46^. Planarian *stx5* is ubiquitously expressed and is required for tissue integrity and survival. Therefore, we suggest that *stx5* may play non-synaptic roles in tissue homeostasis and potentially regeneration.

Importantly, given our criteria of neural expression and altered behavior after inhibition, we identified stx1 and stxbp1-1 as most likely components of the neuronal synapse. To confirm this connection, we wanted to assess whether knocking down *stx1* and *stxbp1-1* impacted synapses themselves. We therefore assessed the density of synaptic connections in the brain using anti-synapsin staining (Fig. 4C). We confirm that knocking down either *stx-1* or *stxbp1-1* caused a decrease in synaptic volume, measured by the neuropil-to-body ratio (Fig. 4D). Additionally, the intensity of synapsin staining was significantly reduced after *stxbp1-1*(RNAi) (Fig. 4E). Because of the high level of similarity between human syntaxin binding protein 1 and planarian Stxbp1-1, we reasoned that an available human antibody could detect the planarian protein. Indeed, anti-syntaxin binding protein 1 labeling marked synaptic spaces including the neuropil and core of the ventral nerve cords, similar to the synapsin staining (Fig. 4F). To confirm the specificity of this antibody, we stained control and knockdown animals with anti-syntaxin binding protein 1. We found that both the area and intensity of staining was reduced, after *stxbp1-1*(RNAi) (Fig. 4G-I). We similarly saw STXBP1-1 staining decreased after *stx1*(RNAi), supporting our argument that the proteins encoded by these genes work together. Taken together, our work shows that *stx1* and *stxbp1-1* are expressed in the nervous system, required for proper movement, and work together to coordinate proper brain regeneration through their role in synapse formation and/or maintenance.

## Discussion

This work identified two potential synapse protein-encoding genes that promote planarian neuronal function—*stx1* and *stxbp1-1*. Our work indicates that STX1 and STXBP1-1 are *bona fide* planarian synapse proteins, which is an important in furthering a foundational understanding of chemical synapses in this critical regenerative model organism. Here we also demonstrated that existing antibodies targeting the human sequence of STXBP1 works in planarians. Using existing antibodies to STX1 and STXBP1-1 from other animals could allow us to better assess synapse formation in different populations of neurons. Our work also lays the foundation for uncovering how synapses reform during regeneration in planarians. Knocking down *stx-1* or *stxbp1-1* resulted in a reduction in brain size due to decreased neuropil volume. This suggests that neurotransmission or vesicle trafficking at the synapse promotes maturation or stability of synaptic structures, which in turn promotes neuron survival. This has been shown in *Xenopus*, *Drosophila*, mice, and humans^47–50^. Additionally, synaptic deficits have been associated with neurodegenerative diseases and aging, which further supports that hypothesis^51,52^. Finally, a critical question for regenerative neuroscientists, including those of us studying planarians, is whether regenerative neurogenesis follows the “use it or lose it” principle common to other model organisms, including rodents, chicks, gerbils, and cats^53–59^. Do planarians create an excess of neurons and only those with successful, active synaptic connections survive? In other organisms, syntaxin and syntaxin binding protein are required for axon growth and neuron survival in the brain^32,33^. This may also be the case for planarians. STX-1 and STXBP1-1 are likely just two of the key players in the process of synapse form and function and future work could outline the types of neurons that use these proteins for synapses, enabling us to understand how synaptogenesis impacts neuronal survival. Our characterization of STX1 and STXBP1-1 in the planarian synapse have broadened our understanding of neuronal communication and consequences to behavior in a highly regenerative bilaterian.

### Experimental Procedures

#### Planarian Care

Planarians used are from an asexual clonal line (*Schmidtea mediterranea* Strain CIW4) kept in a 1X Montjuïc salt solution (1.6 mmol NaCl, 1.0 mmol CaCl_2_, 1.0 mmol MgSO_4_, 0.1 mmol MgCl_2_, 0.1 mmol KCl and 1.2 mmol NaHCO_3_ in Milli-Q water) in a dark container at 18°C^60^. Planarians were fed every two weeks with beef liver paste (White Oak Pastures) and supplemented with 50 µg/mL Gentamicin sulfate (Gemini Bio-Products) once per month. Animals were starved a minimum of 7 days prior to experiments and animals undergoing RNAi were dosed with Gentamicin after feeding and amputation.

#### Identification of genes of interest and cloning

Putative homologs were identified by using BLASTx in PlanMine^61^ using human and mice syntaxin and syntaxin-binding proteins (Supp. Table 1). These candidates were verified via reciprocal BLAST and domain analysis using translated protein queries using InterproScan. 20-22 bp primers were designed to amplify a 700 bp segment of each transcript of interest using Primer3plus^62^. We amplified genes of interest for cloning into EAM1105I-digested pJC53.2 vector with standard molecular methods^63^. Sequence validation and orientation were determined through Sanger sequencing (GENEwiz/Azenta).

#### Phylogeny & domain analysis

Phylogenetic analysis was completed in phylogeny.fr using the top model organism hits from different families of syntaxin and syntaxin-binding proteins^64^. Planarian homologs were named according to phylogenetic position and BLAST similarity to known proteins. Planarian protein domains were identified within a translation of the longest open reading frame with InterProScan^65^. For protein information, see Supp. Table 1.

#### Riboprobe synthesis, in situ hybridization, immunofluorescence

Antisense riboprobes were synthesized from PCR-amplified products using either T3 or SP6 polymerase using standard molecular methods^63^. Colorimetric *in situ* hybridization was completed using Digoxigenin-11-UTP (Roche)-labeled probes. Fluorescent *in situ* hybridization probes were synthesized using either Digoxigenin-11-UTP (Roche) or Fluorescein-12-UTP (Roche). Colorimetric *in situ* hybridization was completed as previously described^66^. Briefly, planarians starved for seven days were placed in a 15 mL conical tube for untreated ISH or small glass vials of each for ISH after RNAi. Planarians were killed with 7.5% N-Acetyl Cysteine in 1X PBS (made from 10X PBS: 1.37 M NaCl, 27 mM KCl, 100 mM Na_2_HPO_4_, 20 mM KH_2_PO_4_, pH 7.4) incubating for 7 minutes. Then, the planarians were fixed in 4% fixative solution (4% formaldehyde in PBSTx (PBS + 0.3% Triton X-100) for 20 minutes. Animals were then washed with PBS before dehydration to methanol for overnight storage at -20°C. Animals were then rehydrated to PBS and bleached (5% formamide, 1.25% H_2_O_2_, 0.5x SSC; 1X SSC = 150 mM NaCl and 15 mM sodium citrate). After further washes, animals were hybridized in riboprobe solution (riboprobe diluted in a solution of 50% de-ionized formamide, 25% 20x SSCx, 100 µg/mL yeast RNA, 1% Tween-20, and 50 mg/mL dextran sulfate in DEPC-treated water), with riboprobes diluted 1:1000 to 1:2000 depending on probe strength. After hybridization at 56°C for 17 hours, the samples were washed in 2X SSC three times and 0.2X SSC four times and incubated in a blocking solution (5% horse serum in TNTx, which is 0.1 M Tris pH 7.5, 0.15 M NaCl, 0.3% Triton-X-100) for 1 to 2 hours. The blocking solution was removed before the samples were incubated in 500 µl of antibody solution (blocking solution with 1:2000 anti-digoxigenin Fab fragments conjugated with alkaline phosphatase, Roche), rocking for 15-18 hours at 4°C. After eight TNTx washes, animals were developed using NBT (Roche) and BCIP (Roche) in alkaline phosphatase buffer (100 mM Tris pH 9.5, 100 mM NaCl, 50 mM MgCl_2_, 0.1% Tween-20, 10% Polyvinyl alcohol) in the dark until they reached the desired staining. To stop the development, planarians were washed in PBSTx for 5 minutes twice, then washed in 500 µl of 100% ethanol for 20 minutes. After another wash of 500 µl of PBSTx for 5 minutes, PBSTx was replaced with an 80% glycerol solution and the planarians were transferred to glass slides for imaging.

For fluorescent ISH, animals were processed using the protocol above until the blocking step. FISH blocking solution instead consisted of 0.5% Roche Western Blocking Reagent and 5% horse serum in TNTx. The antibody solution included either anti-DIG-POD (Roche) or anti-FITC-POD (Roche) at a dilution of 1:2000. Tyramide conjugates for fluorescent development were synthesized as previously described^67^. For double fluorescent ISH, we used DIG- and FITC-labeled probes. After washing off the DIG antibody solution, a 10-minute development was completed with TAMRA-tyramide [1:500 TAMRA-tyramide, 1:1000 4-IPBA, 0.05% H_2_O_2_ in TSA Buffer (2M NaCl in 0.1M Boric acid pH 8.5, filter sterilized)]. After washing, antibody inactivation proceeded with a 45-minute wash in 0.1 M sodium azide in PBSTx, followed by a second blocking step and the FITC-antibody solution. After further washes in TNTx, development was conducted with FAM-tyramide (1:500 FAM-tyramide, 1:1000 4-IPBA, 0.05% H_2_O_2_ in TSA Buffer) for 10 minutes. After further washes, samples were stained overnight in DAPI solution (1 µg DAPI/10 mL TNTx) and then washed 6 times in TNTx. Fluorescent samples were mounted in Vectashield antifade mounting medium (Vector Labs).

Immunofluorescence was completed as previously described^16^. Briefly, animals were killed in 6% HCl, followed by 20 minutes of 4% fixative (4% formaldehyde in PBSTx (PBS + 0.3% Triton X-100) and further washes. Animals were bleached in 6% H_2_O_2_ in PBSTx for 13 hours. After further washes, animals were blocked in 1% bovine serum albumin (BSA) in PBSTx for four hours. Animals were incubated in the primary antibody (anti-synapsin (1:100, 3C11 concentrate; Developmental Studies Hybridoma Bank)^20^ or anti-syntaxin-binding-protein-1 (1:175, Cusabio)) overnight at 4°C for 16 hours. After eight more 20-minute PBSTx washes, animals were blocked again for two hours and then incubated in the secondary antibody overnight at 4°C for 16 hours (goat-anti-mouse Alexa Fluor 488 (1:500, Invitrogen) or goat-anti-rabbit Alexa Fluor 488 (1:200, Invitrogen)). Similar to fluorescent ISH, samples were washed six more times in 30-minute PBSTx washes and then stained overnight in DAPI solution (1 µg DAPI/10 mL TNTx) and then washed 6 times in TNTx. Samples were mounted in Vectashield antifade mounting medium (Vector Labs).

#### Imaging

Live animals and animals from colorimetric ISH were imaged using a Zeiss Axio Zoom V16 microscope and an Axiocam 506 Color camera utilizing ZEN 2.3 pro software. Videos of live animals were taken with an iPhone 12 and processed into still images. Fluorescent imaging was performed on an LSM 880 confocal microscope with an upright AXIO imager Z2 and Zen Black 2.3 SPI software. Images of heads were taken with the 20x/0.8 objective and images for co-expression analysis were captured using the 40x/1.4 objective. Whole body images were taken with the 20x/0.8 objective and tiling. Images were processed using Photoshop and Illustrator (Adobe). Max intensity projections created using Fiji^68^ are shown for whole head images. For cilia imaging, individual animals were placed in a ring of petroleum jelly on a 24x60 mm coverslip within a single drop (roughly 50 µL) of 1X Montjuïc salt solution. A 22x22 mm coverslip was placed on top to form a closed chamber. For high-speed video analysis at 250 fps, we used an inverted Eclipse Ti2 microscope (NIKON) equipped with a long-distance DIC condenser and a 10x 0.95 Planapo objective and 2.5x adapter. Images were recorded using an EoSens 3CL camera (Mikrotron) and a CORE2 DVR Express rapid storage device (IO Industries). Recordings were exported in AVI format and analyzed in Fiji is ImageJ^68^.

#### RNA interference

dsRNA of interest was synthesized from pJC53.2 clones via transcription with T7 RNA polymerase using standard molecular techniques^69^. Animals were separated into 60x15 mm Petri dishes 3 days before feedings with 12 worms 3-4 mm in length per experiment. Planarians were fed 5 μg dsRNA mixed with 1 μl McCormick green food dye and 30 μl beef liver paste. dsRNA targeting *Aequorea victoria green fluorescent protein (GFP)* serves as a negative control. For double knockdowns, planarians were fed 5 µg of each gene’s dsRNA for a total of 10 µg per feeding. The control group received 10 µg of *GFP* dsRNA, and single knockdowns received 5 µg of the dsRNA of interest along with 5 µg of *GFP* to control feeding amounts. The double knockdown received 5 µg each of *syntaxin-1* dsRNA and *syntaxin-binding protein 1-1* dsRNA (Supp. Fig. 2). For all RNAi experiments, animals were allowed to eat for 2-4 hours before being washed and transferred to a new Petri dish where they were treated with Gentamycin sulfate at a dilution of 1:1000 (50 mg/mL stock from Gemini Bio-Products). Feeding paradigms are described in relevant figures.

#### Behavioral assays

Movement Assays: Three animals from each condition were pipetted into the center of a test plate, a 60x15 mm Petri dish in 10 mL of 1X Montjuïc salt solution over a printed circle 1.5 cm in diameter. After 1 minute, the position of the animals, inside or outside of the circle, was recorded. Videos were also taken of live animals moving in a 60x15 mm Petri dish, with snapshots every five seconds showing relative animal position.

Righting Assay: Each animal was pipetted into a 60x15 mm Petri dish and placed ventral side up. The time each animal took to flip dorsal side up was recorded, with righting completed when the entire length of the worm was dorsal side up. Animals that could not flip within six minutes were counted as not having flipped during the experimental duration.

Stimuli Response Assays: Animals were considered to have responded to stimuli if they contracted their body or moved away from the stimulus. For touch stimulus, an eyebrow hair was used to touch the trunk of an animal and its immediate response was noted. For visual stimulus, a blue laser pointer (XIMBI) was pointed at the trunk of an animal and its immediate response was recorded.

#### Image Quantification and Statistical Analysis

All images and videos were blinded before quantification. Brain-to-body and neuropil-to-body ratios were traced in triplicate^70^. Co-expression was calculated by randomly selecting 25 cells per image expressing a gene of interest and identifying the proportion that express the other marker.

Animal speed was quantified by importing videos into Fiji (Fiji is Image J^68^) and using the FFMPEG plugin. Minimum intensity projections were used to trace the trajectory of individual animals. The length traveled over time was used to determine speed, as described previously^71^. Kymograms were generated from videos using the KymoResliceWide plugin^72^ in FIJI^68^. Beat frequency was calculated by analyzing peaks present over time, as described previously^73^. Cilia coordination was categorized by the degree of vertical synergy between different cilia. We assessed a minor defect in coordination when we saw large groups of cilia moving in sync in groups with at least two different patterns. Kymograms with gaps in motile cilia (spaces without lines) were categorized as uncoordinated. For categorical data, individual pairwise comparisons between control(RNAi) and knockdowns were completed using Fisher’s exact test, two-sided; to correct for multiple comparisons, a two-stage linear step-up procedure of Benjamini, Krieger, and Yekutieli^74^ was performed with Q = 5%, estimated number of null hypotheses 2 to provide the adjusted p-value, which has been reported. Fluorescence intensity was measured by calculating the average intensity per animal (maximum value of fluorescence at 5 different regions of interest in Z space) and subtracting the average background (fluorescence measurement of unstained background of three areas per image). All statistical analyses were completed using GraphPad Prism version 10.0.0 or 11.0.0 for Mac OS X, GraphPad Software, Boston, Massachusetts. Bar graphs show mean and standard deviation as well as individual data points. Statistical test information is also included in figure legends.

## Acknowledgements

We would like to thank past and current Roberts-Galbraith lab members who assisted in project feedback and support. We would like to thank Dr. Muthugapatti Kandasamy and the Microscopy core for training and technical support. We would also like to thank Dr. Nikolay Filipov for assistance with data acquisition and analysis. This study was completed through funding from the National Institute of Neurological Disorders and Stroke (NINDS) of the National Institutes of Health under award number 5R01NS128096. RRG is also supported by the Lois K. Miller Professorship at the University of Georgia (UGA). KFL was supported by the National Institute of General Medical Sciences of the National Institute of Health under award number R35GM152057. KBC was supported by the National Institute of General Medical Sciences of the National Institute of Health under award number 1T32GM142623 and NINDS project numbers 5R25NS107179 and F31NS134272. KBC was supported by the University of Georgia Research Foundation and ARCS Foundation. PAR was supported by funding from the Center for Undergraduate Research Opportunities and the Office of Experiential Learning at UGA. This content is solely the responsibility of the authors and does not represent the official views of funding agencies.

---

**Supplemental Figure 1:**
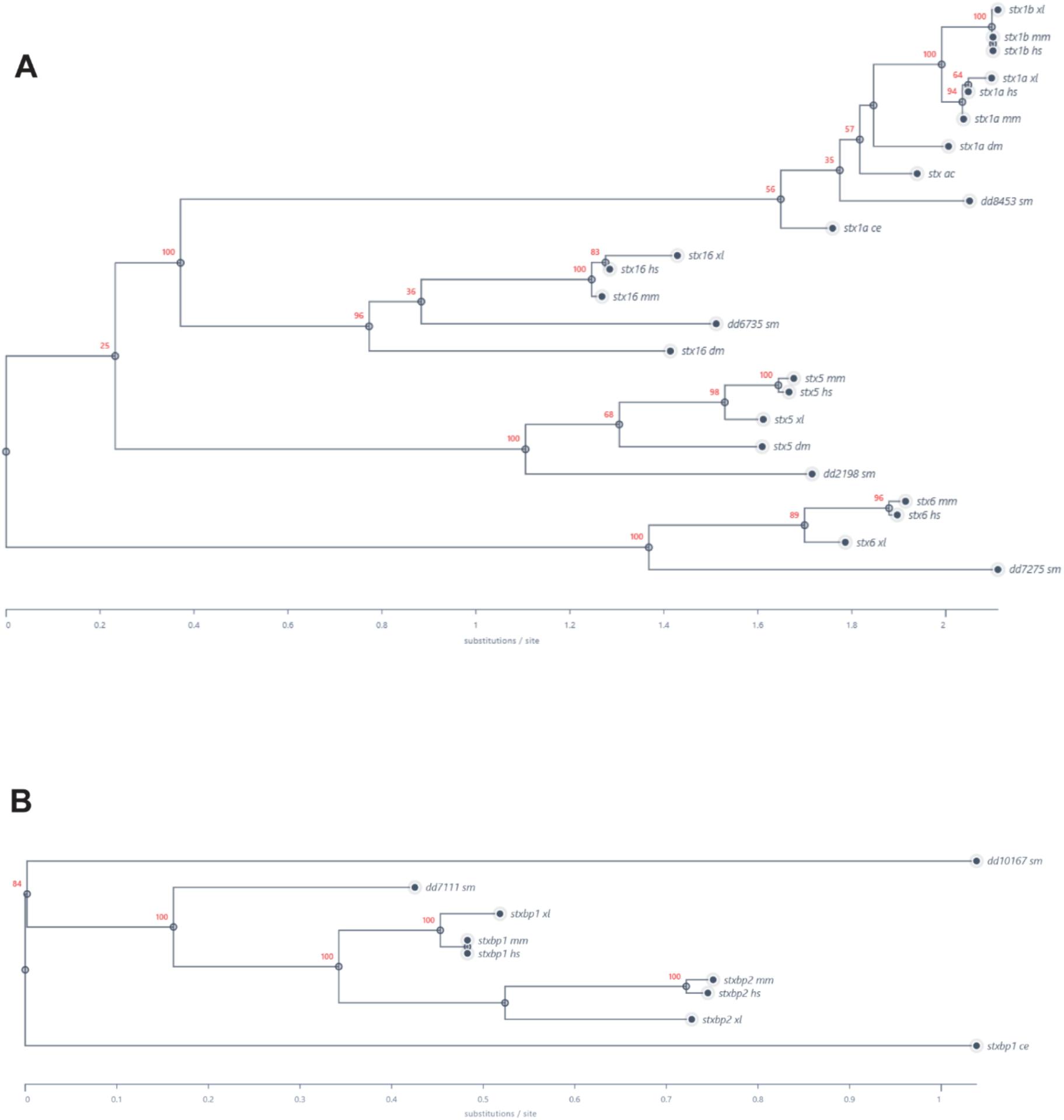
Phylogeny of planarian STX and STXBP places them in distinct families. A) Syntaxin protein phylogeny. B) Syntaxin-binding protein phyogeny. Genus and species initials are shown after the underscore for each protein. *Homo sapiens* (hs), *Mus musculus* (mm), *Xenopus laevis* (xl), *Caenorhabditis elegans* (ce), *Drosophila melanogaster* (dm), *Schmidtea mediterranea* (sm). Red numbers indicate the percentage of branch support for each node.

**Supplemental Figure 2:**
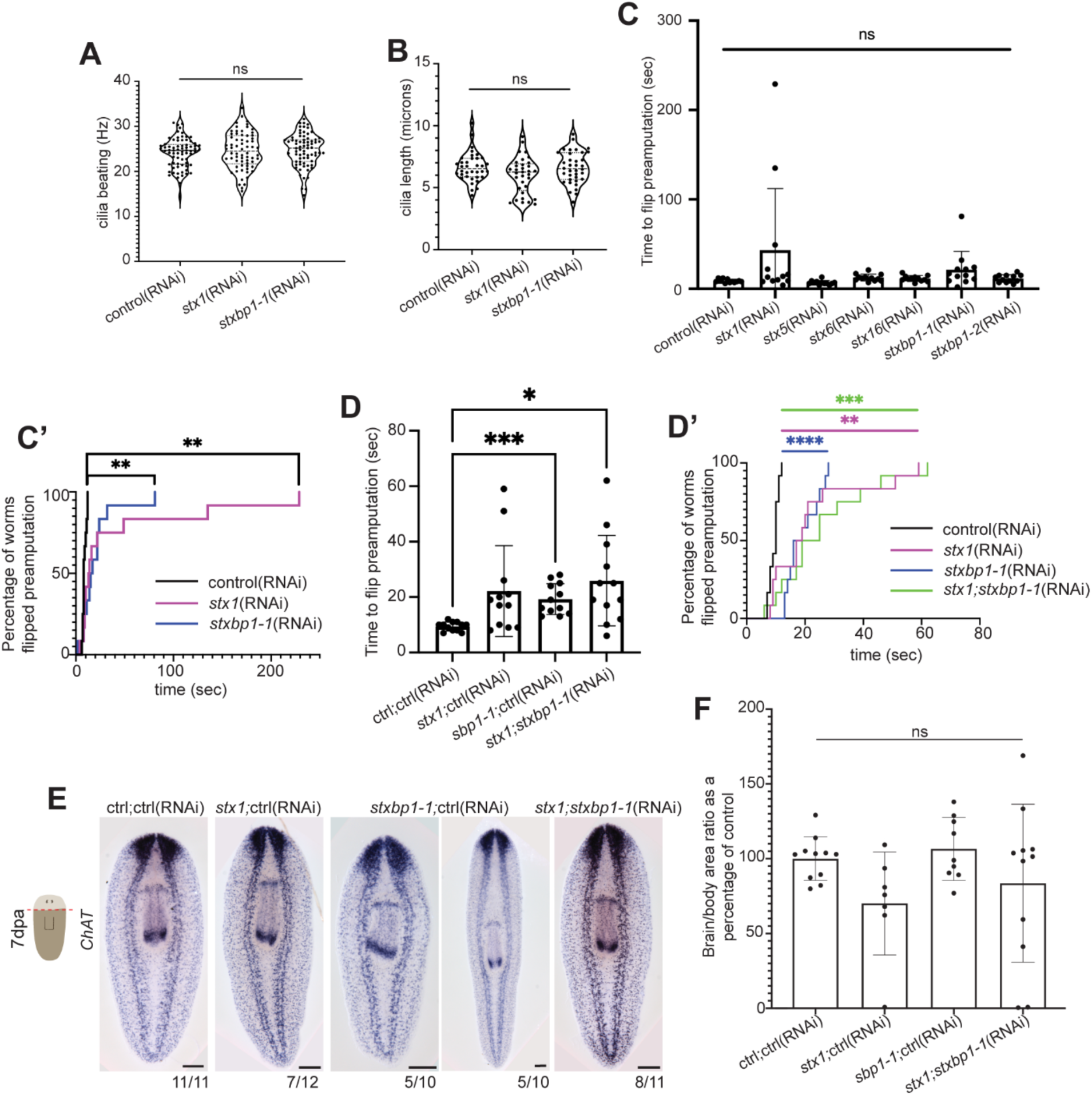
Additional information on *stx1* and *stxbp1-1 RNAi phenotypes*. A-B) Quantification of cilia beat frequency and length, respectively, with significance assessed using one-way ANOVA with Dunnett’s multiple comparisons test. n = 6 measurements/animal; N=11 animals for *stx1*(RNAi) and 12 animals each for control(RNAi) and *stxbp1-1*(RNAi). C) Righting assay depicting average time to flip pre-amputation. One-way ANOVA with Dunnett’s multiple comparisons test. A’) Percentage of animals flipped over time, with significance assessed using a Mantel-Cox log rank test. D-D’) Righting assay depicting time to flip in double knockdown animals pre-amputation. D) One-way ANOVA with Dunnett’s multiple comparisons test. D’) Mantel Cox log-rank test. E) Colorimetric ISH depicting brain regeneration marked by *ChAT* in double knockdowns. F) Quantification of E. One-way ANOVA with Dunnett’s multiple comparisons test. Scale bar = 200 µm (E). ns = not significant, * P ≤0.05, ** P ≤ 0.01, *** P ≤0.001, **** P ≤ 0.0001.

**Supplemental Figure 3:**
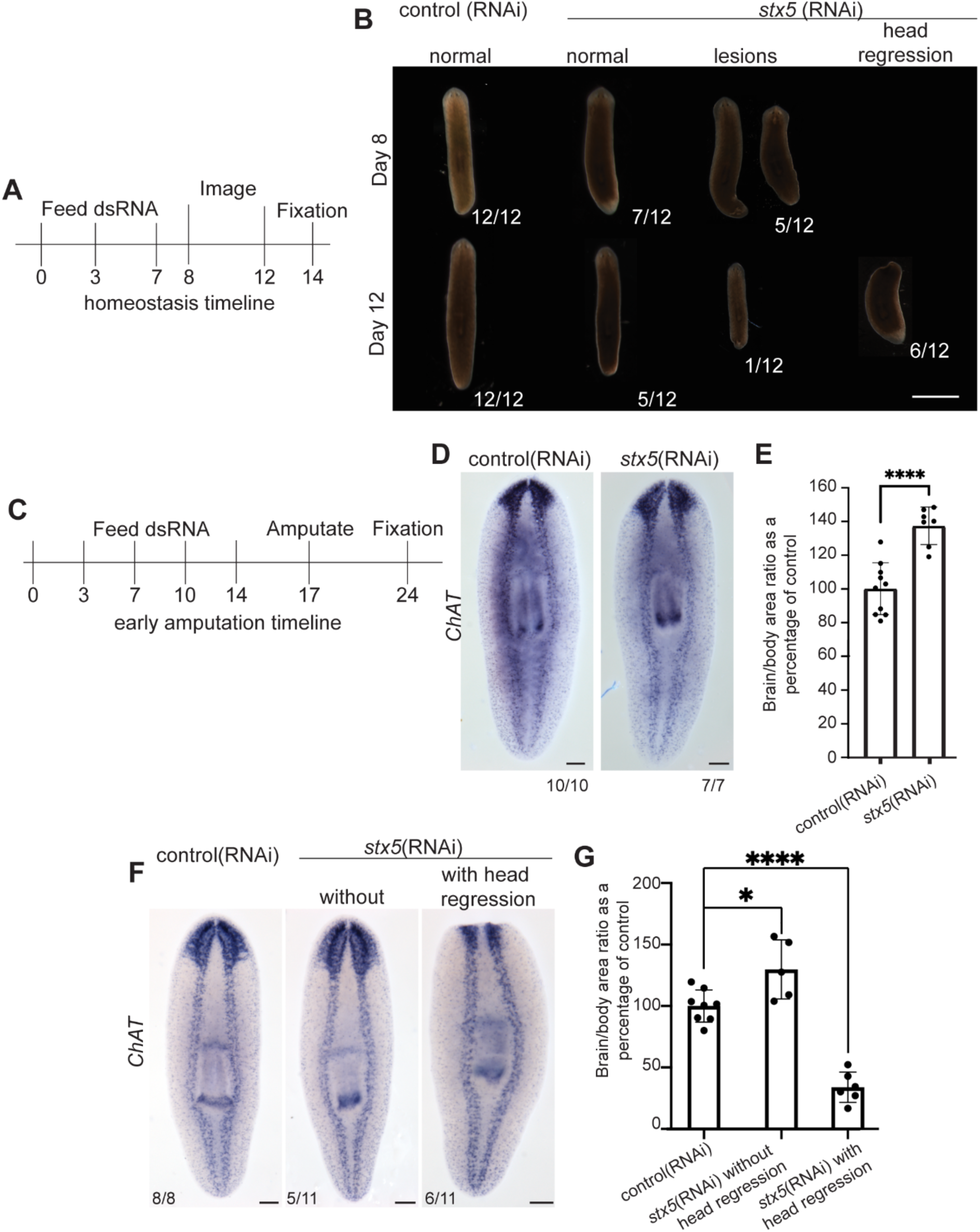
*stx5* causes lesions, lysis, and larger brains in homeostasis and regeneration. A) Homeostasis timeline. B) Morphology of control(RNAi) or *stx5*(RNAi) over time depicting instances of lesions and head regression. C) Early amputation timeline. D) Colorimetric ISH depicting brain regeneration marked by *ChAT*. E) Quantification of D, with significance assessed using an unpaired t-test. F) Colorimetric ISH depicting brain size in the absence of injury marked by *ChAT*. G) Quantification of F. One-way ANOVA with Dunnett’s multiple comparisons test. Scale bars = 200 µm (B, F), 2000 µm (D). ns = not significant, * P < 0.05, ** P < 0.01, *** P < 0.001, **** P < 0.0001.

**Supplemental Table: Primer and Protein Sequences.** Tab 1) Planarian primer sequences for cloning and protein sequences used for homology search. Tab 2) Syntaxin and syntaxin binding protein sequences from other organisms used for phylogeny and domain analysis. Tab 3) BLASTx results and transcriptional atlas expression for planarian genes. Tab 4) tBLASTn search of human syntaxin and syntaxin binding proteins to identify the closest planarian homolog.

